# Analysis of Treacher Collins syndrome 4-associated mutations in *Schizosaccharomyces pombe*

**DOI:** 10.1101/2025.04.13.648503

**Authors:** Kei Kawakami, Hiroaki Kato

## Abstract

Treacher Collins syndrome (TCS) is a rare congenital disorder characterized by craniofacial deformities. Although mutations in several genes involved in ribosome biogenesis have been identified in patients with TCS, the molecular mechanisms by which these mutations exert their effects remain poorly understood. In this study, we established *Schizosaccharomyces pombe* models for TCS type 4 (TCS4) by introducing specific mutations in the *rpa2* gene, which encodes Rpa2, the second largest subunit of RNA polymerase I (Pol I). TCS4-associated *rpa2*^R1022C^ and *rpa2*^R1022S^ mutations impaired cell growth without altering Rpa2 protein levels. These mutants were defective in 35S pre-rRNA biogenesis and were sensitive to the Pol I inhibitor BMH-21. These findings underscore the essential role of Rpa2 residues associated with TCS4 in rRNA transcription and cell growth.

## Introduction

Treacher Collins syndrome (TCS) is a rare congenital disorder characterized by craniofacial deformities resulting from mutations in several genes involved in ribosome biogenesis, including TCOF1, POLR1C, POLR1D, and POLR1B [1]. Type 4 TCS (TCS4) specifically involves Arg1003Cys, Arg1003Ser, or Ser682Arg mutations in POLR1B, which encodes RPA2, the second largest subunit of RNA polymerase I (Pol I) [2, 3]. The high degree of conservation of these amino acid sequences from yeast to humans underscores their fundamental roles in the function of RPA2 [2]. Investigating the effects of these mutations on Pol I transcription at the cellular level is crucial for understanding this disease and developing therapeutic strategies. However, the functional consequences of these mutations on Pol I activity remain unclear.

The fission yeast *Schizosaccharomyces pombe* has been instrumental in elucidating fundamental biological processes because of its genetic tractability and the conservation of key cellular pathways with higher eukaryotes [4]. In *S. pombe*, ribosomal RNAs (rRNAs) are encoded by a ribosomal DNA array located near both ends of chromosome III. These rRNA genes are transcribed by Pol I within the nucleolus as a polycistronic precursor (i.e., the 35S pre-rRNA) [5]. The 35S pre-rRNA undergoes sequential processing at the external transcribed spacers (5′ ETS and 3′ ETS) and internal transcribed spacers (ITS1 and ITS2) to produce the mature 18S, 5.8S, and 28S rRNAs [6]. In *S. pombe*, the Pol I complex comprises 14 subunits. Seven of these subunits—Nuc1, Rpa2, Rpa49, Rpa43, Rpa34, Ker1, and Rpa12—are specific to Pol I. Nuc1 and Rpa2 are the largest subunits forming the catalytic core for rRNA transcription [7–10]. Rpa49, Rpa34, Ker1, and Rpa12 are nonessential subunits that facilitate transcriptional initiation, elongation, and termination [11–17].

Due to its high similarity to human Pol I in both subunit composition and three-dimensional structure [10, 18–20], *S. pombe* is expected to serve as a valuable model for investigating human diseases associated with Pol I dysfunction.

In this study, we established genetically tractable models to investigate the pathophysiology of TCS4 by introducing analogous mutations into *S. pombe rpa2*, which is homologous to POLR1B. We examined the phenotypic effects of these mutations on *S. pombe* cell growth and rRNA transcription. Through these analyses, we uncovered the cellular and molecular defects caused by TCS4 mutations, shedding light on the molecular defects involved in the pathogenesis of TCS4.

## Materials and Methods

### *S. pombe* strains and media

The *S. pombe* strains used in this study were derived from the standard strain L972 (Table S1). *S. pombe* media were prepared as described previously [21].

### Site-directed mutagenesis of the genomic *rpa2* ORF

Cells harboring the *rpa2*^R1022C/S^ mutation were generated using a PCR-based method with long primers containing site-directed mutations [22, 23]. Integration and base substitution were confirmed using colony PCR and Sanger sequencing, respectively. The detailed strategy is shown in the Supplementary Material (Fig. S1).

### Spot assay

Cells were grown to the stationary phase in YES medium at 30°C. Five-fold serial dilutions were prepared, spotted onto plates, and incubated for 3 days before imaging. For the drug sensitivity analysis, YES plates containing 300 μM BMH-21 (S7718; Selleck Biotech, Kanagawa, Japan) were used.

### Protein extraction and western blotting

Protein extraction and western blotting were performed as previously described [9]. Cells were cultured to log phase in YES medium at 30°C. Briefly, 5 × 10^7^ cells were harvested and re-suspended in 150 μL of buffer 1 (50 mM HEPES, 140 mM NaCl, 1 mM EDTA, 1% TritonX-100, 0.1% sodium deoxycholate, pH 7.5). Subsequently, 150 μL of 2× Laemmli buffer was added, and the cell suspension was heated at 95°C for 5 min. Zirconia beads (11079105z; Biospec Products, OK, USA) were added to the cell suspension, followed by cell lysis (MULTI BEADS SHOCKER; YASUI KIKAI, Osaka, Japan) for 10 min at 2,700 rpm. To completely denature the proteins, the samples were boiled again at 95°C for 5 min. For western blotting, anti-FLAG (F3165, M2; Sigma-Aldrich, St. Louis, MO, USA) and anti-β-tubulin (63-160; Bioacademia, Osaka, Japan) were used as primary antibodies. HRP-conjugated anti-mouse IgG (W4021; Promega, Madison, WI, USA) and HRP-conjugated anti-rabbit IgG (W4011; Promega, Madison, WI, USA) were used as secondary antibodies. The signals were visualized using ImageQuant 800 (Cytiva, Tokyo, Japan).

### Construction of template plasmids for northern probe

DNA fragments corresponding to 494 bp of 28S rRNA, 403 bp of 5′ ETS, and 177 bp of 5S rRNA were amplified by PCR using oligonucleotide primers KNB-269 and KNB-270, KNB-289 and KNB-290, and KNB-275 and KNB-276, respectively. The amplified fragments were cloned into the BamHI site, downstream of the T7 RNA polymerase promoter in the pGEM-3Z vector (P2151; Promega, Madison, WI, USA). The constructs were confirmed by Sanger sequencing. The primers and plasmids used for probe construction are listed in Tables S2 and S3, respectively.

### Preparation of RNA probes for northern blotting

Template plasmids were linearized with HindIII (1060A; TAKARA Bio, Shiga, Japan) and subjected to in vitro transcription using T7 RNA polymerase (2540A; TAKARA Bio).

### Northern blotting

Total RNA was extracted as described previously [24]. Total RNA (~0.5 μg) was separated on a 1% denaturing agarose gel. After electrophoresis, the RNA was transferred onto a nylon membrane (Hybond-N^+^) (RPN1210B; Cytiva, Tokyo, Japan) using 10× SSC and crosslinked by UV irradiation. Hybridization and DIG-based signal detection were performed using the DIG Northern Starter Kit (12039672910; Roche, Basel, Switzerland). The signals were visualized using ImageQuant 800 (Cytiva, Tokyo, Japan).

## Results

### Amino acid residue implicated in TCS4 is evolutionarily conserved

To generate an *S. pombe* model for TCS4, we first examined the conservation of the amino acids related to TCS4 between *H. sapiens* RPA2 and *S. pombe* Rpa2. As previously reported, alignment analysis revealed that the arginine at position 1003 of human RPA2 was conserved at position 1022 in *S. pombe* Rpa2 (Fig. 1A and 1B) [2, 10]. In the human Pol I complex, R1003 is located at the cleft, a hybrid-binding domain close to the nascent RNA and template DNA strands in the Pol I complex (Fig. 1A and 1C) [18–20]. We compared the spatial arrangement of R1003 in *H. sapiens* RPA2 and R1022 in *S. pombe* Rpa2 within the reported Pol I elongation complexes [10, 19]. Based on this comparison, R1022 of Rpa2 is located in the cleft region and adopts a spatial arrangement similar to that of R1003 in human RPA2 (Fig. 1C). Thus, *S. pombe* is considered an excellent model organism for analyzing TCS4 mutations.

**Fig. 1.**
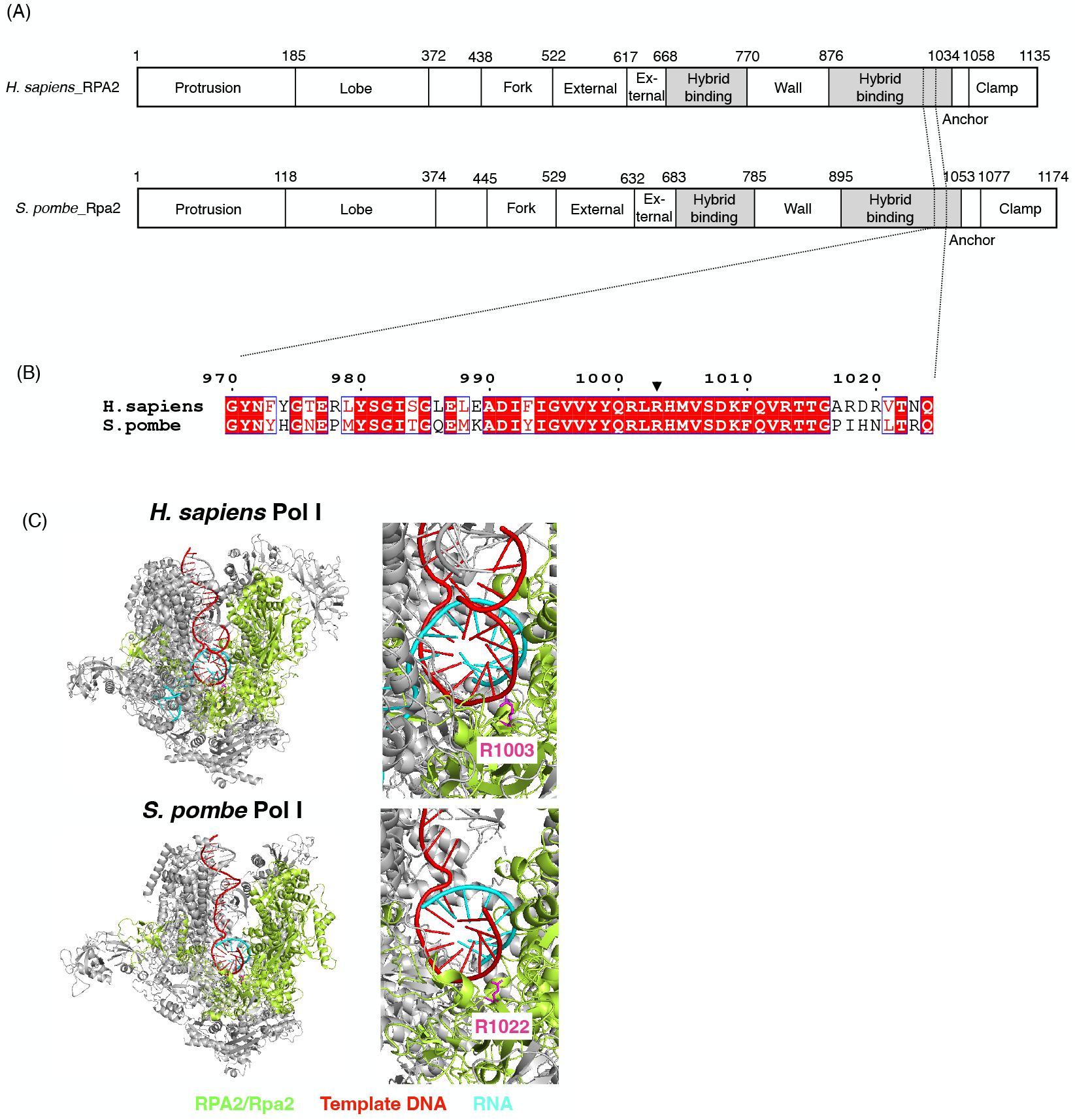
Amino acid residue implicated in TCS4 is evolutionarily conserved. (A) Schematic domain architecture of the *H. sapiens* RPA2 (NP_061887) and *S. pombe* Rpa2 (NP_595819) [10, 18]. (B) Alignment analysis of a part of the hybrid-binding domain in *H. sapiens* RPA2 and corresponding residues in *S. pombe* Rpa2. The amino acid residues conserved among *H. sapiens* and *S. pombe* are indicated with white letters on a red background. Arrowhead shows TCS4-associated residue R1003 in *H. sapiens* and R1022 in *S. pombe*. Alignment was performed using ClustalW and visualized with ESPript 3.0 [28]. (C) Structural comparison of the Pol I elongation complexes between *H. sapiens* (PDB: 7OB9, upper) and *S. pombe* (PDB: 7AOE, lower), based on the structures reported by Misiaszek et al., 2021 (DOI: 10.1038/s41594-021-00693-4) and Heiss et al., 2021 (DOI: 10.1038/s41467-021-21031-8), respectively [10,19]. Enlarged images of the cleft region are shown on the right. RPA2/Rpa2, template DNA, and RNA are shown in green, red, and cyan, respectively. R1003 of *H. sapiens* RPA2 and R1022 of *S. pombe* Rpa2 are highlighted in magenta. This figure is reproduced under the terms of the Creative Commons Attribution 4.0 International License (CC BY 4.0).

### Generation of *S. pombe* model for TCS4

To investigate the effects of TCS4-associated mutations in *S. pombe, rpa2*^R1022C^ or *rpa2*^R1022S^ mutations were introduced into haploid cells (Fig. S1). Sanger sequencing confirmed the successful introduction of the mutations into the *rpa2* ORF (Fig. 2A). To examine the effect of mutations on cell growth, the growth rates were assessed in YES liquid medium at 30°C. As a result, the *rpa2*^R1022C^ and *rpa2*^R1022S^ mutants were found to exhibit growth defects compared to the wild-type (Fig. 2B). To determine whether the *rpa2*^R1022C/S^ mutation affected Rpa2 protein levels, western blot analysis was performed. The results showed that the protein levels of Rpa2^R1022C/S^-5FLAG were comparable to those of wild-type Rpa2-5FLAG (Fig. 2C). These findings suggest that the *rpa2*^R1022C/S^ mutations impair cell growth without altering Rpa2 protein levels.

**Fig. 2.**
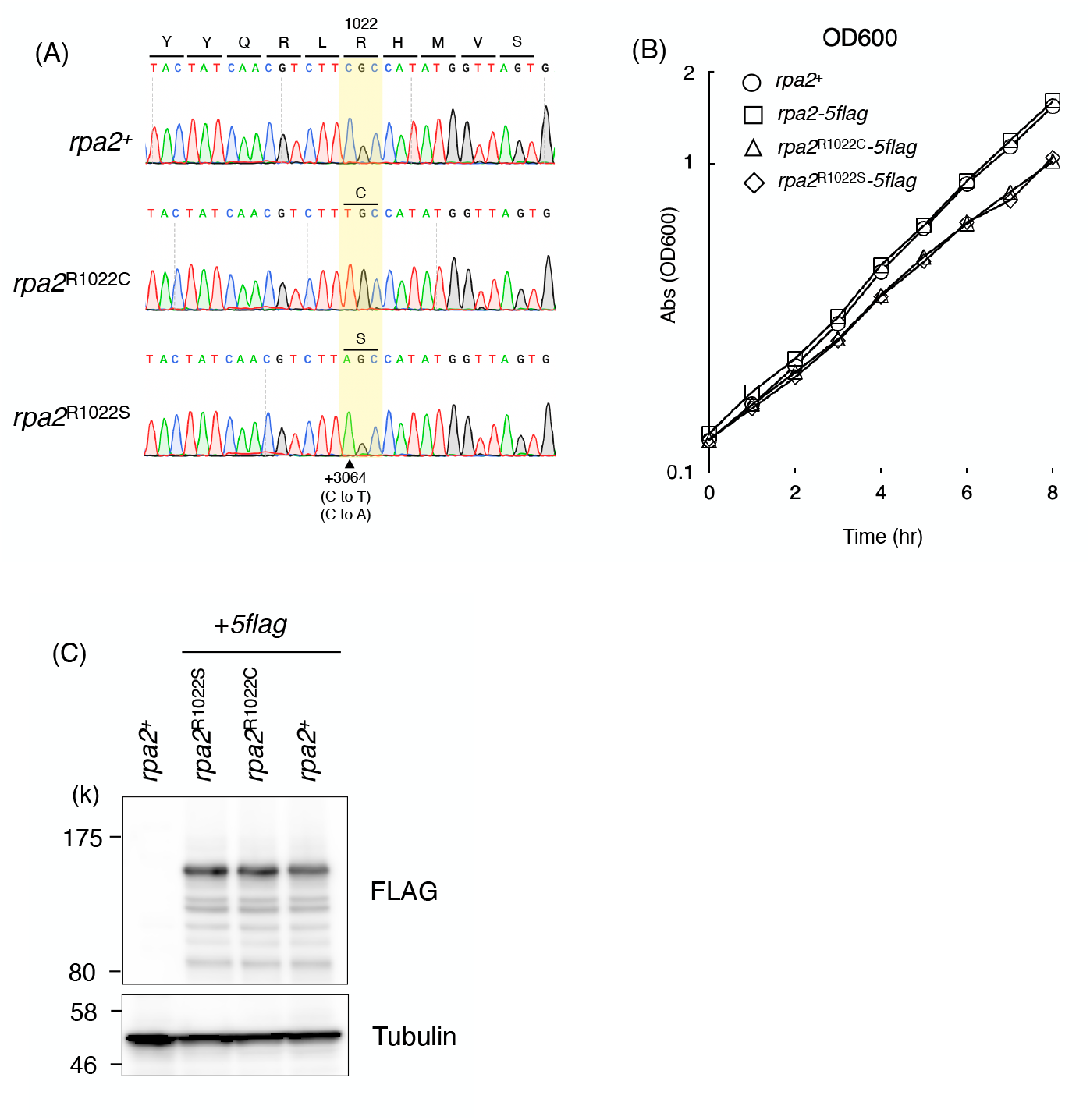
Generation of *S. pombe* model for TCS4. (A) The sequences from +3049 to +3079 bp of the *rpa2* ORF in wild-type, *rpa2*^R1022C^, and *rpa2*^R1022S^ are shown. The base substitution C3064T or C3064A resulted in the R1022C or R1022S missense mutation, respectively. (B) Analysis of growth rate in YES medium. Wild-type, *rpa2*^R1022C^, and *rpa2*^R1022S^ cells were grown in YES liquid medium at 30°C. The absorbance at 600 nm of each culture in the log phase was measured every hour and plotted. The vertical axis represents the OD600 absorbance on a logarithmic scale, while the horizontal axis indicates the cultivation time. (C) Western blotting analysis of Rpa2-5FLAG proteins. Total proteins extracted from non-tagged, wild-type, *rpa2*^R1022C^, and *rpa2*^R1022S^ cells were analyzed with anti-FLAG. Tubulin was analyzed for the loading control.

### Rpa2 R1022C/S mutations impair rRNA transcription

The 28S, 18S, and 5S rRNAs are derived from a single 35S pre-rRNA transcript via multiple processing events. To assess the impact of the *rpa2*^R1022C/S^ mutations on 35S pre-rRNA biogenesis, northern blot analysis was performed using an RNA probe targeting the 5′ETS region, which is specific for 35S pre-rRNA [6]. A significant reduction in 35S pre-rRNA levels was observed in the *rpa2*^R1022C^ and *rpa2*^R1022S^ mutants (Fig. 3A and 3B), suggesting that rRNA transcription efficiency was markedly reduced in these mutants. Consistent with this, *rpa2*^R1022C^ and *rpa2*^R1022S^ mutants exhibited sensitivity to the Pol I-specific inhibitor BMH-21 (Fig. 3C) [25]. Interestingly, the levels of mature 28S rRNAs in *rpa2*^R1022C^ and *rpa2*^R1022S^ mutants were comparable to those in the wild-type (Fig. 3A and 3B). These findings indicate that *rpa2*^R1022C/S^ mutations cause defects in 35S pre-rRNA biogenesis.

**Fig. 3.**
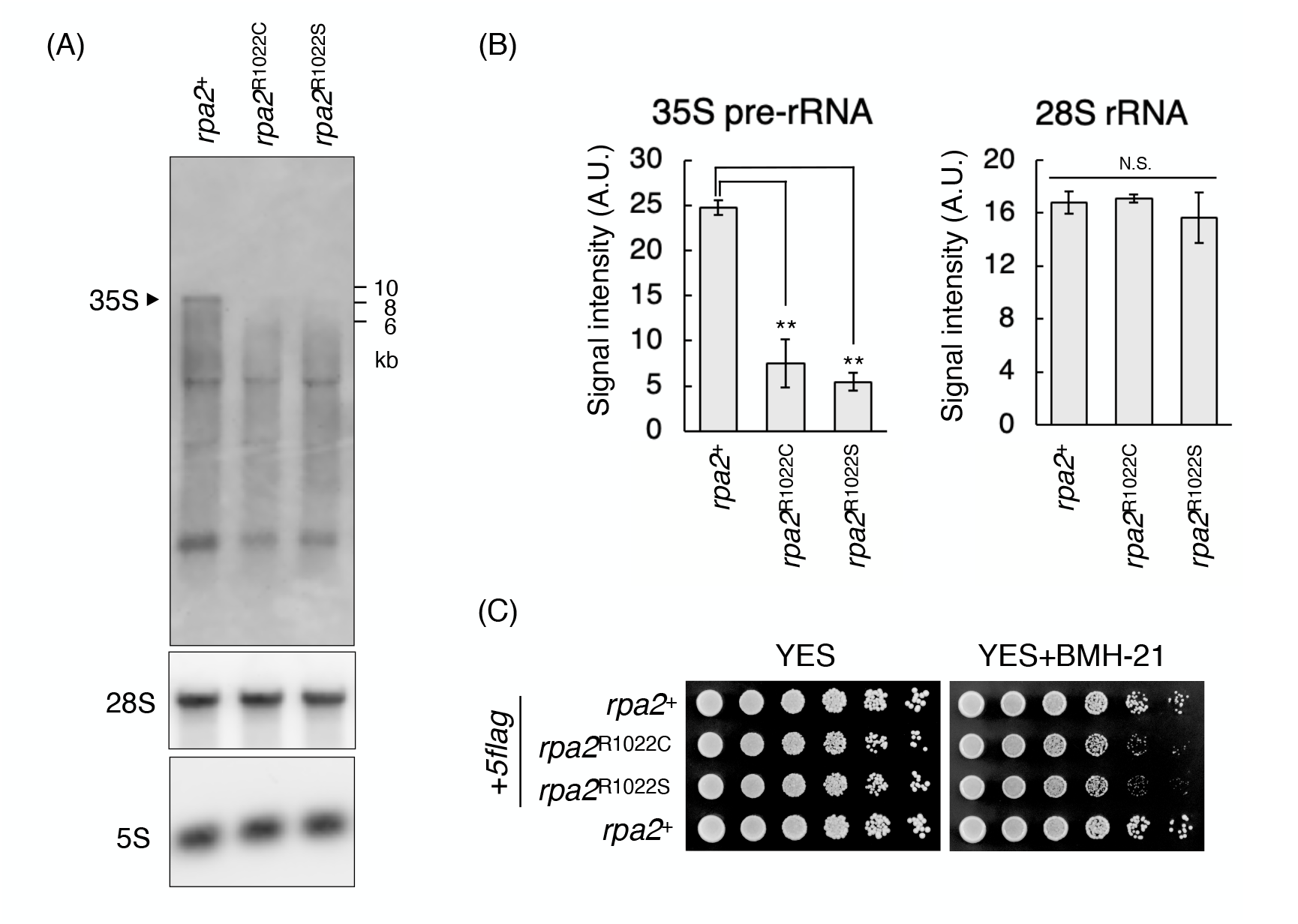
TCS4-associated mutations affect 35S pre-rRNA biogenesis. (A) Northern blotting analysis of 35S pre-rRNAs and 28S rRNAs in indicated strains. The image of 5S rRNA is shown as a loading control. (B) Quantitative analysis of 35S pre-rRNAs and 28S rRNAs is shown in (A). The signal intensities of the bands corresponding to 35S pre-rRNAs or 28S rRNAs were quantified using ImageJ. Each signal intensities were normalized to those of 5S rRNAs. The vertical axis represents the signal intensities in arbitrary units. Error bars show the standard deviation from three independent samples. Statistical significance was determined by Student’s t-test. (\*\**p* < 0.01, N. S. = not significant). (C) Spot assay for testing BMH-21 sensitivity. Five serial dilutions of indicated cell suspension were spotted on YES and YES + 300 μM BMH-21 plates and incubated at 30°C for 3 days.

## Discussion

In this study, a model system was established to investigate the molecular basis of TCS4 in *S. pombe*. Structural studies of human Pol I have suggested that R1003 of RPA2 is located in a hybrid-binding domain within the cleft where it stabilizes DNA and/or DNA/RNA hybrids within the polymerase complex to ensure proper RNA synthesis (Fig. 1A and 1C) [18–20]. However, the effects of TCS4 mutations on Pol I transcription have not been reported. Our data clearly showed that 35S pre-rRNA transcription was impaired by *rpa2*^R1022C/S^ mutation (Fig. 3A and 3B), suggesting that R1022 is a critical residue for efficient transcription. Generally, arginine carries a positive charge and interacts with the negatively charged phosphate backbone of DNA via electrostatic interactions. In *Pyrococcus furiosus* RNA polymerase, arginine residues in the cleft are essential for binding to the template and non-template DNA strands to promote transcriptional elongation [26]. The loss of arginine-mediated interactions due to R1022C/S mutations may destabilize the transcription bubble, leading to slower transcriptional elongation or an increased frequency of transcriptional pausing and premature termination. Further biochemical studies are needed to elucidate how these mutations affect nucleic acid binding or the protein-protein interactions involved in Pol I transcription.

Despite the reduction in the amount of 35S pre-rRNA, the mature 28S rRNA levels in *rpa2*^R1022C/S^ mutants were comparable to those in the wild-type strain (Fig. 3A and 3B). These findings suggest that the half-life of mature rRNAs can be dynamically regulated within cells to store enough amount of mature rRNAs. Post-transcriptional modifications stabilize the secondary structure of rRNA and prevent their degradation [27]. Interestingly, the *rpa2*^R1022C/S^ mutants exhibited growth inhibition, despite having mature rRNA levels similar to those of the wild-type strain (Fig. 2B, 3A, and 3B). A similar phenotype has been reported in other Pol I-related mutants [17]. The functionality of mature rRNA in these mutants remains unclear, and further analysis of the ribosome-mediated protein synthesis efficiency is needed to clarify this point. Future studies using *rpa2*^R1022C/S^ mutants should focus on elucidating the molecular mechanisms underlying TCS and other ribosomopathies.

## Author contributions

Conceptualization, K. K.; methodology, K. K. and H. K.; investigation, K. K.; resources, K. K. and H. K.; writing of the original draft, K. K.; reviewing and editing, H. K.; funding acquisition, K. K. and H. K.

## Data availability

All the data obtained in this study are included in the main paper and Supplementary Material.

## Declaration of generative AI and AI-assisted technologies in the writing process

During the preparation of this work, the authors used ChatGPT to improve the readability and language of the manuscript. After using this tool, the authors reviewed and edited the content as required and took full responsibility for the content of the published article.

## Acknowledgments

We thank Dr. E. Obayashi for his valuable support and helpful discussions. We also thank the Interdisciplinary Center for Science Research, Shimane University, for the use of their facilities.

