## supplementary_material for "Analysis of Treacher Collins syndrome 4-associated mutations in *Schizosaccharomyces pombe*"

**Table S1.** Strains used in this study

| Strain ID | Genotype |
|---|---|
| L972 | *h*$^{-}$ |
| IZM86 | *h*$^{-}$ *rpa2-5flag*::*kanMX6* |
| IZM85 | *h*$^{-}$ *rpa2*$^{R1022C}$*-5flag*::*kanMX6* |
| IZM93 | *h*$^{-}$ *rpa2*$^{R1022S}$*-5flag*::*kanMX6* |
| Bio9308 | *h*$^{-}$ *leu1-32 ura4-D18 rpa2-5flag*::*kanMX6* |

**Table S2.** Primers used in this study

| Oligo ID | Sequence (5′ → 3′) | Description |
|---|---|---|
| KNB-129 | GTTTCATGTGTCCAGTTCAC | *rpa2* fw |
| KNB-133 | GAGTTGTCTACTATCAACGTCTTTGCCATATGGTTAGTGATAAATTTC | R1022C (C3064T) |
| KNB-134 | GAAATTTATCACTAACCATATGGCAAAGACGTTGATAGTAGACAACTC | R1022C (G3064A) |
| KNB-135 | GAGTTGTCTACTATCAACGTCTTAGCCATATGGTTAGTGATAAATTTC | R1022S (C3064A) |
| KNB-136 | GAAATTTATCACTAACCATATGGCTAAGACGTTGATAGTAGACAACTC | R1022S (G3064T) |
| KNB-137 | CATACGAATGTGTGCTCACG | *rpa2* rv |
| KNB-289 | CCCGGATCCGGTGGTTGAAAGGAGAAAAG | *5'ETS* |
| KNB-290 | CCCGGATCCCCACTTCTTTTTCCCCTTCT | *5'ETS* |
| KNB-269 | CCCGGATCCGAGACCGATAGCGAACAAGTAG | *28S rRNA* |
| KNB-270 | CCCGGATCCGGGTCCCAACAGCTATGCTC | *28S rRNA* |
| KNB-275 | CCCGGATCCGTCTACGGCCATACCTAGG | *5S rRNA* |
| KNB-276 | CCCGGATCCTTTGATGTCGAAAACGAGGA | *5S rRNA* |

Red characters: Mismatch mutation to produce R1022C/S; blue characters: BamHI site

**Table S3.** Plasmids used in this study

| Plasmid ID | Description |
|---|---|
| pKK_6 | pGEM-3Z_28SrRNA |
| pKK_9 | pGEM-3Z_5SrRNA |
| pKK_11 | pGEM-3Z_rRNA5'ETS |

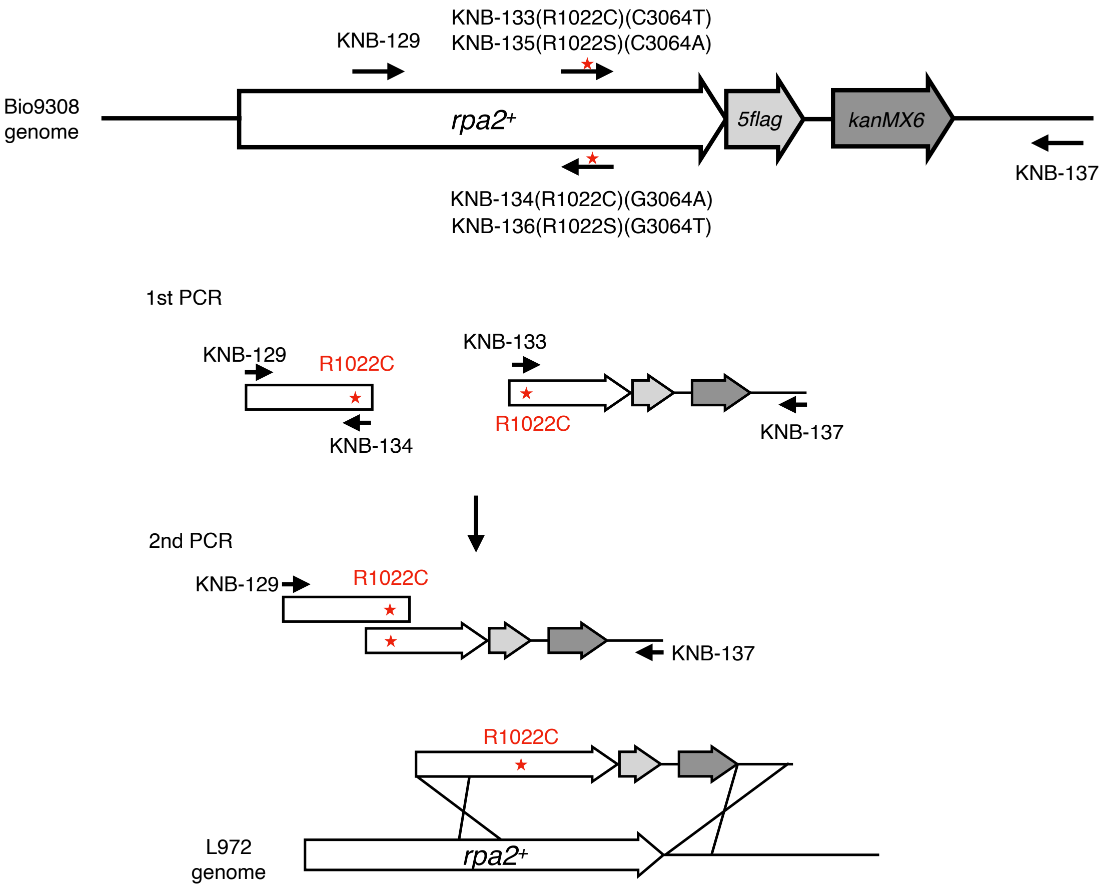


**Fig. S1. Site-directed mutagenesis of the genomic *rpa2* ORF**

A DNA fragment containing the R1022C mutation was generated using a two-step PCR procedure. In the first step, genomic DNA derived from the Bio9308 strain (*rpa2-5flag*::*kanMX6*) was used as the template. PCR was performed using the primers KNB-129 and KNB-134 or KNB-133 and KNB-137 to generate two DNA fragments: one containing the R1022C mutation and the other containing R1022C with *5flag*::*kanMX6*. In the second step, these two fragments were joined by PCR using the primers KNB-129 and KNB-137. This yielded a *rpa2*$^{R1022C}$-*5flag*::*kanMX6* fragment with homologous sequences on the C-terminus of *rpa2*, which was used to transform the wild-type strain L972. For the R1022S mutation, primers KNB-129 and KNB-136 or KNB-135 and KNB-137 were used in the first PCR step. All other procedures were identical to those used to generate R1022C.

Uncropped western blot images (Related to Fig. 2C)

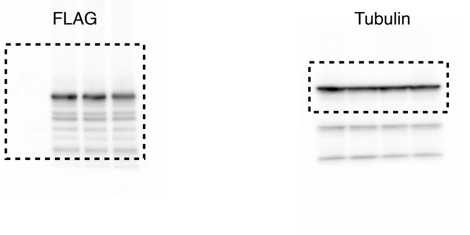


Uncropped northern blot images (Related to Fig. 3A)

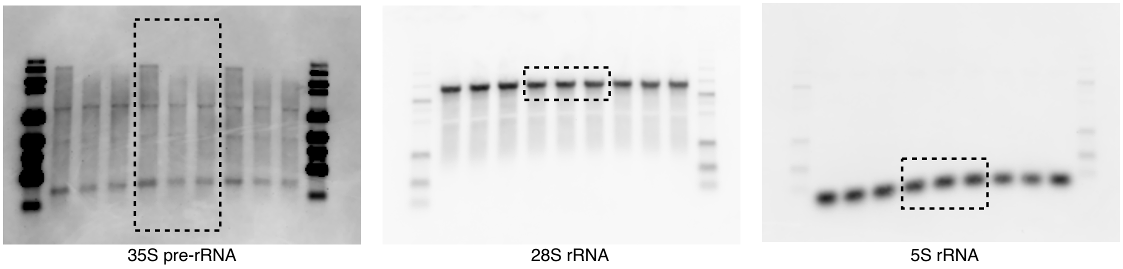


**Fig. S2. Uncropped blotting images**

Uncropped western (upper panel) and northern (lower panel) blots are shown. Areas surrounded by dashed lines are used in the main figures.
